# Metabolic and Redox Pathway Dysregulation in HIV-Associated Coronary Endothelial Dysfunction

**DOI:** 10.1101/2025.11.26.690743

**Authors:** Kshipra S. Keole, Syed Bukhari, Anum Minhas, Charles D. Cohen, Amelia Wallace, Damani Piggott, Thorsten M. Leucker, Kevin Sun, Luigi Adamo, Allison Hays

## Abstract

**Background:** People with HIV (PWH), even with sustained viral suppression on antiretroviral therapy (ART), remain at increased risk for cardiovascular disease. Coronary endothelial dysfunction, a sensitive marker of early vascular injury and a potential target for intervention is common in this population, but its biological basis remains unknown.

**Methods:** We performed a cross-sectional study combining *in vivo* coronary MRI and high-throughput serum proteomics to investigate mechanisms of coronary endothelial dysfunction in treated HIV. Forty-five virally suppressed PWH and twenty-nine age- and sex-matched healthy controls underwent coronary MRI during isometric handgrip exercise to quantify coronary endothelial function, defined as the percentage change in coronary cross-sectional area (%CSA) from rest to stress. An increase in coronary CSA <2% indicated endothelial dysfunction. Parallel serum proteomic profiling was performed using the SomaScan 7K platform, and differential protein expression between groups was analyzed using linear modeling (LIMMA).

**Results:** Coronary endothelial dysfunction was more prevalent in PWH with suppressed viral load compared to controls (67% vs 10%, p<0.001). Pathway analysis of differentially expressed proteins between participants with and without endothelial dysfunction highlighted significant dysregulation of glutathione dependent detoxification, oxidative metabolism and fatty acid β-oxidation pathways in individuals with endothelial dysfunction (adjusted p value <0.05).

**Conclusions:** Endothelial dysfunction in PWH on ART is associated with metabolic and redox imbalance. These findings highlight glutathione and fatty acid oxidation related pathways as potential therapeutic targets for reducing cardiovascular risk in this patient population.

## INTRODUCTION

With contemporary antiretroviral therapy (ART), people with HIV (PWH) are living longer, yet have a persistently elevated burden of coronary artery disease (CAD) that remains a leading cause of morbidity and mortality in this patient population (1,2).

The pathophysiology of this excess risk of CAD is complex. Cardiometabolic risk factors like diabetes, dyslipidemia and hypertension are prevalent in PWH and can be worsened by certain antiretroviral medicines (3). Chronic infection and ART are associated with metabolic derangements and a redox imbalanced environment that predisposes to vascular disease, even when viral replication is suppressed (4).

More specifically, several studies have demonstrated impaired mitochondrial oxidative phosphorylation and increased reliance on glycolytic metabolism in circulating endothelial and immune cells in PWH, indicating a state of metabolic dysregulation (5). This shift is accompanied by accumulation of lipid intermediates, reduced fatty acid β-oxidation and increased reactive oxygen species (ROS) generation (4,6). ART-associated dyslipidemia which is characterized by elevated triglycerides and reduced high density lipoprotein (HDL), further contributes to vascular inflammation and oxidative stress (7,8). Together, these interrelated metabolic and inflammatory disturbances are associated with a proatherogenic vascular milieu, linking chronic HIV infection to persistent endothelial injury and heightened cardiovascular risk.

Impaired coronary endothelial function (CEF) plays a central role in the initiation and progression of atherosclerosis. It predicts future events and is frequently present in PWH even in the absence of obstructive CAD (9,10). Therefore, CEF serves not only as a sensitive marker of subclinical vascular injury but also as a promising therapeutic target for early intervention to mitigate downstream cardiovascular risk.

Prior work from our group and others, linked HIV-related metabolic perturbations, including elevated proprotein convertase subtilisin/kexin type 9 (PCSK9), to impaired nitric oxide-mediated CEF (11–13). Despite these advances, critical gaps remain; the biological basis of endothelial dysfunction in virally suppressed HIV remains unexplored.

To address this gap, we integrated *in vivo* coronary MRI based measures of CEF during isometric handgrip stress, an established endothelium dependent stimulus, with high-throughput serum proteomic profiling (SomaScan 7K) in PWH on stable ART and matched controls. Proteomic profiling offers an unbiased, high-throughput approach to characterize the circulating protein landscape, enabling investigation of the pathophysiological basis of endothelial dysfunction. We hypothesized that proteomic signatures associated with impaired endothelial function would reveal abnormal metabolic and inflammatory pathways contributing to vascular dysfunction. Our primary goal was to identify molecular drivers of impaired CEF in PWH and people without HIV (PWoH), as well as identify mechanistic targets for reducing residual cardiovascular risk.

## METHODS

### Study Participants

In this prospective cross-sectional study, PWH and age- and sex-matched PWoH were recruited from outpatient clinics at Johns Hopkins Medicine between 2020 and 2025. Eligible participants were ≥21 years old, had no contraindications to MRI, and provided written informed consent under a protocol approved by the Johns Hopkins Medicine Institutional Review Board. PWH were required to be on stable ART for at least one year, have a most recent HIV viral load <200 copies/mL and mean CD4+ cell count >200mm^3^, and have no prior history of myocardial infarction or coronary revascularization. PWoH were confirmed to be HIV-negative. Participants under 50 years of age were required to have no clinical evidence of CAD, whereas those aged 50 or older needed either a coronary artery calcium score of 0 or no luminal disease on prior CT angiography.

### Cardiac Magnetic Resonance Imaging Protocol and Analysis

Cardiac magnetic resonance imaging (CMR) was performed using a commercial 3.0 Tesla (T) whole-body scanner (Achieva, Philips, Best, The Netherlands) equipped with a 32-channel cardiac coil for signal acquisition. Imaging parameters have been described in detail in prior publications (14,15). All participants fasted for at least 8 hours and underwent CMR in the morning before taking any vasoactive medications. Coronary images were obtained perpendicular to a segment of a native coronary artery that had not previously been treated or, if prior intervention was absent, showed no significant stenosis on invasive angiography. To achieve accurate slice alignment perpendicular to the vessel, double-oblique scout scans were performed. Alternating anatomical and velocity-encoded images were acquired both at rest and during 4-7 minutes of continuous isometric handgrip exercise (IHE) at 30% of each participant’s maximal grip strength, using a CMR-compatible dynamometer (Stoelting, Wood Dale, Illinois, USA) under the guidance of a research nurse.

Endothelial function measurements were targeted at a proximal or mid-segment of the coronary artery.

Coronary cross-sectional area (CSA) was quantified from both resting and isometric IHE stress images, and the resulting percentage change (%CSA) was calculated to assess CEF following established protocols. Analyses were conducted independently by two investigators who were blinded to participants’ clinical information. CSA was determined using semiautomated software (Cine Version 3.15.17, General Electric; FLOW Version 4.0, Medis) (16,17). Abnormal CEF was defined as %CSA of less than 2%, corresponding to more than one standard deviation below the mean endothelial response reported in healthy participants from prior studies (18).

### Protein measurements

The relative concentration of plasma proteins or protein complexes in blood collected during patient visits was measured using SOMASCAN, an aptamer-based capture array. The serum was collected using a standardized protocol and sent to SomaLogic for quantification. Approximately 7000 modified aptamers (SOMAmers) were used to measure relative protein concentration.

### SomaLogic quality control

SomaScan quality control and normalization was performed according to the company’s protocols and guidelines.

### Statistical analysis

Principal Component Analysis (PCA) was performed to visualize the degree of separation in the proteomic profiles of the groups of interest. The centroid of each cluster was determined, and the ellipsoid was drawn at 1x standard deviation of the centroid. One visually determined outlier was excluded to improve cluster balance and visual clarity in the normal endothelial function vs abnormal endothelial function comparison. The differentially expressed proteins were identified using the R package limma (linear models for microarray and RNA-seq data) (19). The Benjamini and Hochberg false-discovery rate procedure was used to adjust for multiple comparisons. The two tailed student t-test and the chi-square test were used to compare continuous and categorical variables respectively.

### Bioinformatics analysis

The proteins that were differentially expressed (eg, fold-change ≥1.5, q < 0.05) between participant groups were compiled and analyzed using the ENRICHR databases (20,21). KEGG (Kyoto Encyclopedia of Genes and Genomes), Gene Ontology (GO) Molecular Function, Reactome and Jensen Diseases Curated databases were chosen to evaluate the enrichment of biological pathways.

## RESULTS

### Participant characteristics

We analyzed a cohort of 74 participants that included 45 PWH in complete remission (treated with High Activity Anti-Retroviral Therapy) and 29 matched PWoH. These participants underwent *in vivo* MRI assessment of endothelial function and serum proteomic analysis via Somascan 7K. The demographics of the study population are reported in Table 1 and Supplementary Table 1.

**Table 1.** Baseline Characteristics of Participants by HIV Status

| Variable |  | HIV- (n = 29) | HIV+ (n = 45) | p-value |
| --- | --- | --- | --- | --- |
| Age, years (mean ± SD) | | 52.4 ± 19.4 | 56.9 ± 12.4 | **0.23** |
| Female sex, % |  | 34 | 49 | **0.22** |
| Race, % | **Black** | 45 | 62 | **0.14** |
|  | **White** | 34 | 31 | **0.76** |
|  | **Asian** | 10 | 2 | **0.13** |
| Ethnicity, % | **Hispanic** | 10 | 7 | **0.62** |
|  | **Non-Hispanic** | 90 | 91 |  |
| BMI, kg/m<sup>2</sup> (mean ± SD) | | 27.0 ± 7.6 | 29.3 ± 4.8 | **0.12** |
| Hypertension, % |  | 71 | 62 | **0.56** |
| Diabetes, % |  | 14 | 7 | **0.35** |
| Smoking history, % |  | 25 | 40 | **0.20** |
| Smoking, packs/day (mean ± SD) | | 0.79 ± 0.62 | 1.02 ± 0.79 | **0.49** |
| History substance abuse, % |  | 26 | 43 | **0.15** |
| Total cholesterol, mg/dL (mean ± SD) | | 176.0 ± 37.3 | 183.6 ± 42.3 | **0.45** |
| LDL cholesterol, mg/dL (mean ± SD) | | 96.4 ± 32.4 | 108.1 ± 39.2 | **0.20** |
| HDL cholesterol, mg/dL (mean ± SD) | | 60.7 ± 18.9 | 53.8 ± 15.4 | **0.096** |
| Triglycerides, mg/dL (mean ± SD) | | 98.0 ± 71.3 | 115.7 ± 52.6 | **0.23** |
| Abnormal coronary endothelial function, % |  | 10 | 67 | **<0.001** |
| % Change in Coronary Cross-Sectional Area (mean ± SD) | | 4.28 ± 16.99 | -2.99 ± 10.99 | **0.037** |

Abbreviations: SD = standard deviation; BMI = body mass index; LDL = low-density lipoprotein; HDL = high-density lipoprotein.

We first compared the serum proteome of PWH and PWoH. Supplementary Figure 1 shows that the serum proteome of these two participant groups was almost superimposable when analyzed by PCA (Supplementary Figure 1A). Direct comparison between PWH and PWoH showed few proteins that were differentially expressed (Supplementary Figure 1B). Pathway analysis of these differentially expressed proteins did not highlight any dysregulated biological pathways to a statistically significant level (adjusted p value <0.05). Overall, this suggests that serum proteomic differences between participants with HIV in remission and matched controls were minimal and, if present, were below the statistical threshold of detection of our cohort.

Having completed this important first step, we assessed CEF via cardiac MRI. We measured the %CSA in response to isometric handgrip exercise, a nitric-oxide mediated endothelial-dependent stressor (17). As expected, a greater proportion of PWH exhibited impaired CEF when compared to PWoH (Figure 1A). Approximately two-thirds of HIV-positive participants demonstrated impaired endothelial responses, whereas most control subjects showed preserved function (Figure 1A, χ² = 0.025). The participants were further divided into two groups: “Abnormal endothelial function” and “Normal endothelial function” based on the %CSA values (<2% and ≥2% respectively). Figure 1B shows the specific values of %CSA for all participants in the study, and highlights that most PWH had a markedly impaired coronary vasoreactivity (two-tailed Student’s t test HIV vs non-HIV participants = 6.65 × 10⁻¹⁶).

**Figure 1:**
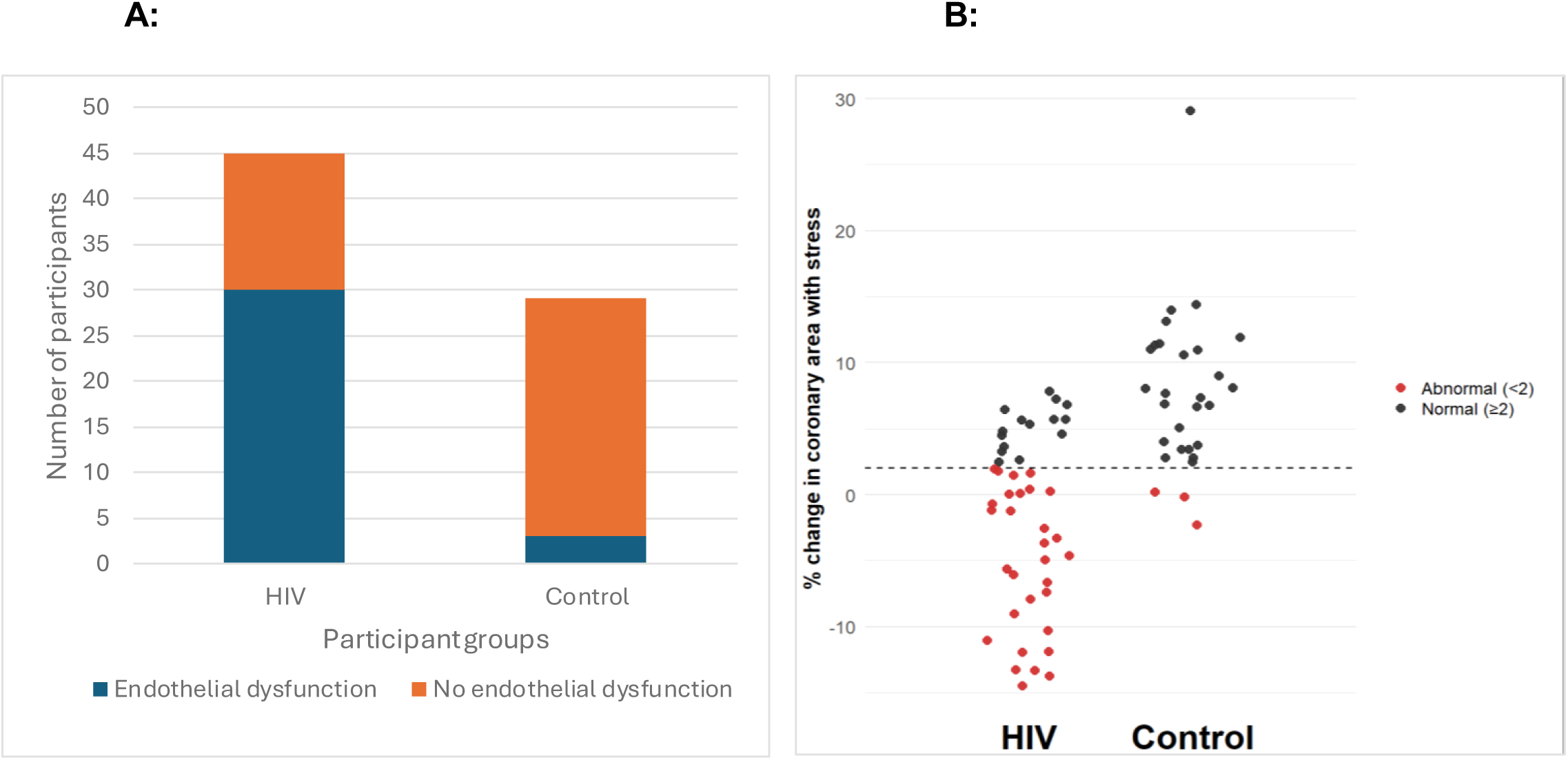
Endothelial dysfunction in participants with HIV and controls. **A)** Prevalence of endothelial dysfunction in HIV-positive and Control participants. The bar graph shows the number of participants with and without endothelial dysfunction in both groups. A higher proportion of endothelial dysfunction was observed in the HIV group compared to the control group (Chi-square p = 0.025). **B)** Coronary endothelial function in HIV and control groups. The scatter plot depicts the percentage change in coronary cross-section area (%CSA) with stress for individual participants in the HIV (left) and control (right) groups. Each point represents a subject, with black dots indicating normal endothelial function (≥2% change) and red dots indicating abnormal endothelial function (<2% change). The dashed horizontal line denotes the 2% threshold for normal endothelial response. Most controls had a normal increase in coronary area, while a higher proportion of HIV-positive individuals exhibited abnormal endothelial function. Two tailed student t test p = 6.65 × 10⁻¹⁶.

To ensure that the observed findings were not driven by major differences in baseline characteristics across groups, we compared demographic and clinical features among participants stratified by HIV infection (Table 1) and by endothelial function (Supplementary Table 2). As shown in Supplementary Table 2, the groups were generally well balanced with respect to age, sex, BMI, and comorbidities commonly associated with endothelial dysfunction, including hypertension, diabetes, and smoking history.

### Serum proteomic profiling of participants with abnormal and normal endothelial function highlights distinctive proteomic signatures

To investigate the biological pathways associated with endothelial function, serum protein expression profiles from participants with normal and abnormal endothelial responses (independent of HIV status) were analyzed using principal component analysis (PCA). The two-dimensional PCAs of the abnormal endothelial function and normal endothelial function groups showed overlapping but distinct clustering of the proteome (Figure 2A). To further investigate this, we performed differential protein expression analysis using limma. Figure 2B reports a volcano plot depicting the log₂ fold change versus -log₁₀(p-value) for each protein, with red and blue points indicating up- and down-regulated proteins, respectively. Proteins surpassing the unadjusted p < 0.05 cutoff were labelled. The most prominent hits included upregulated proteins such as SIRPA, HSP1A1, and GSTT1, and downregulated proteins such as CEBPG, ERAP1, and PILRA.

**Figure 2:**
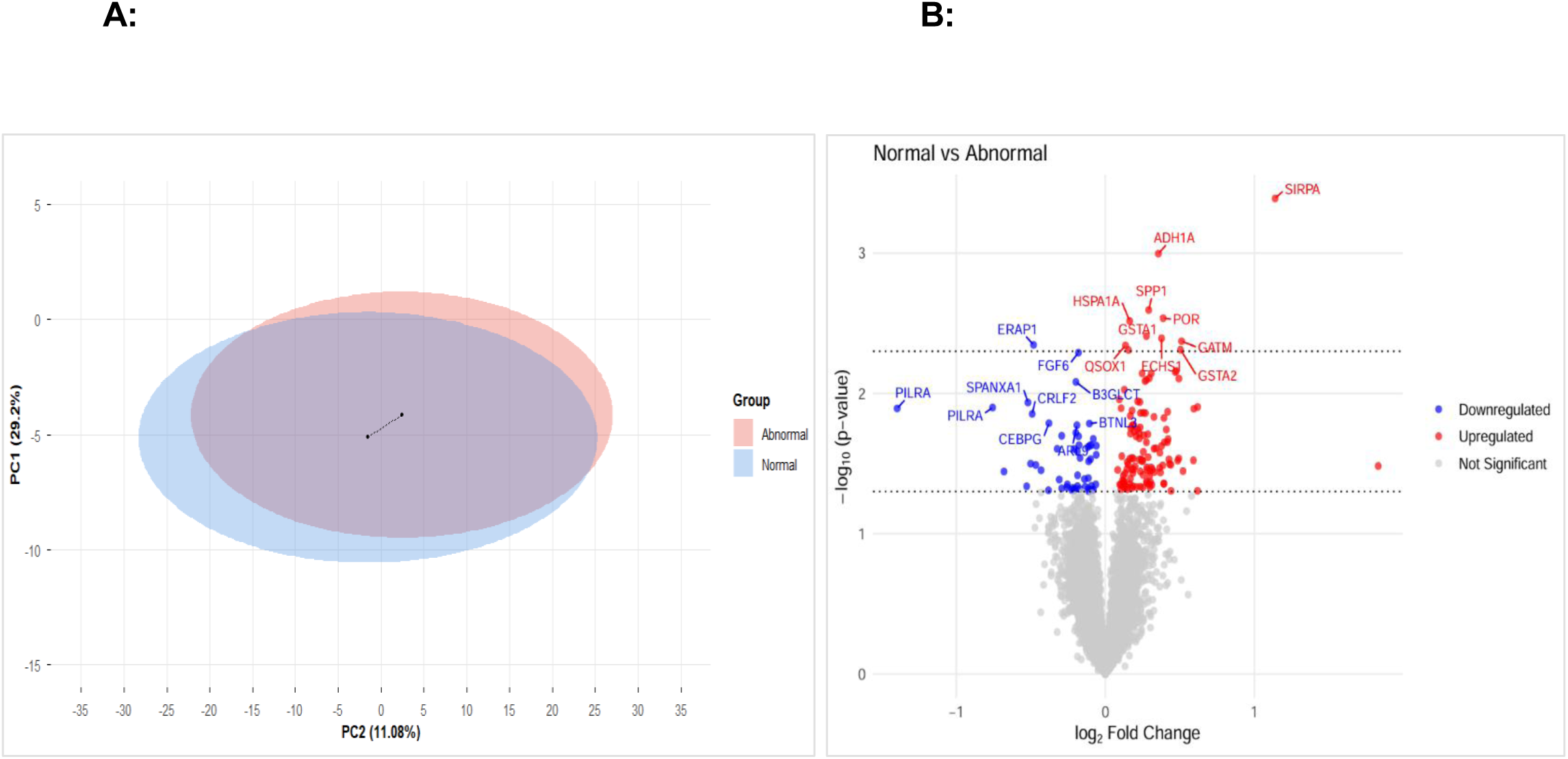
S**e**rum **proteomic analysis of participants with and without endothelial dysfunction A)** 2-dimensional PCA. Endothelial dysfunction is measured according to the values of %CSA with stress. The PCA shows the distribution of individuals with normal (blue) and abnormal (red) endothelial function along the first two principal components. The centroids were calculated based on the PCA vectors. There is partial overlap between the groups, indicating some separation in their underlying profiles but also substantial shared variance. **B)** The volcano plot represents the differential protein expression between participants with normal and abnormal endothelial function. The volcano plot displays log₂ Fold Change (x-axis) versus −log₁₀(p-value) (y-axis) for each protein. Proteins with an unadjusted p value of less than 0.05 are labelled red (upregulated) or blue (downregulated). The black dotted horizontal lines denote unadjusted p-value thresholds of 0.05 and 0.005, respectively. The top 10 upregulated and downregulated proteins are labeled. All proteins with differential expression between the two groups with unadjusted p value < 0.005 were further analyzed via Pathway Analysis.

### Pathways analysis indicates enrichment in glutathione and metabolism related pathways

To evaluate the biological relevance of the differentially expressed proteins, we performed pathway analyses using Enrichr, a web-based tool for comprehensive pathway enrichment analysis (Figure 3). To increase the specificity of our analysis we analyzed proteins differentially expressed with p value <0.005. A total of 12 proteins were submitted for enrichment analysis across four major databases: KEGG, Gene Ontology (GO) Molecular Function, Reactome, and Jensen Diseases Curated. Pathways with an adjusted p < 0.05 were considered statistically significant.

**Figure 3:**
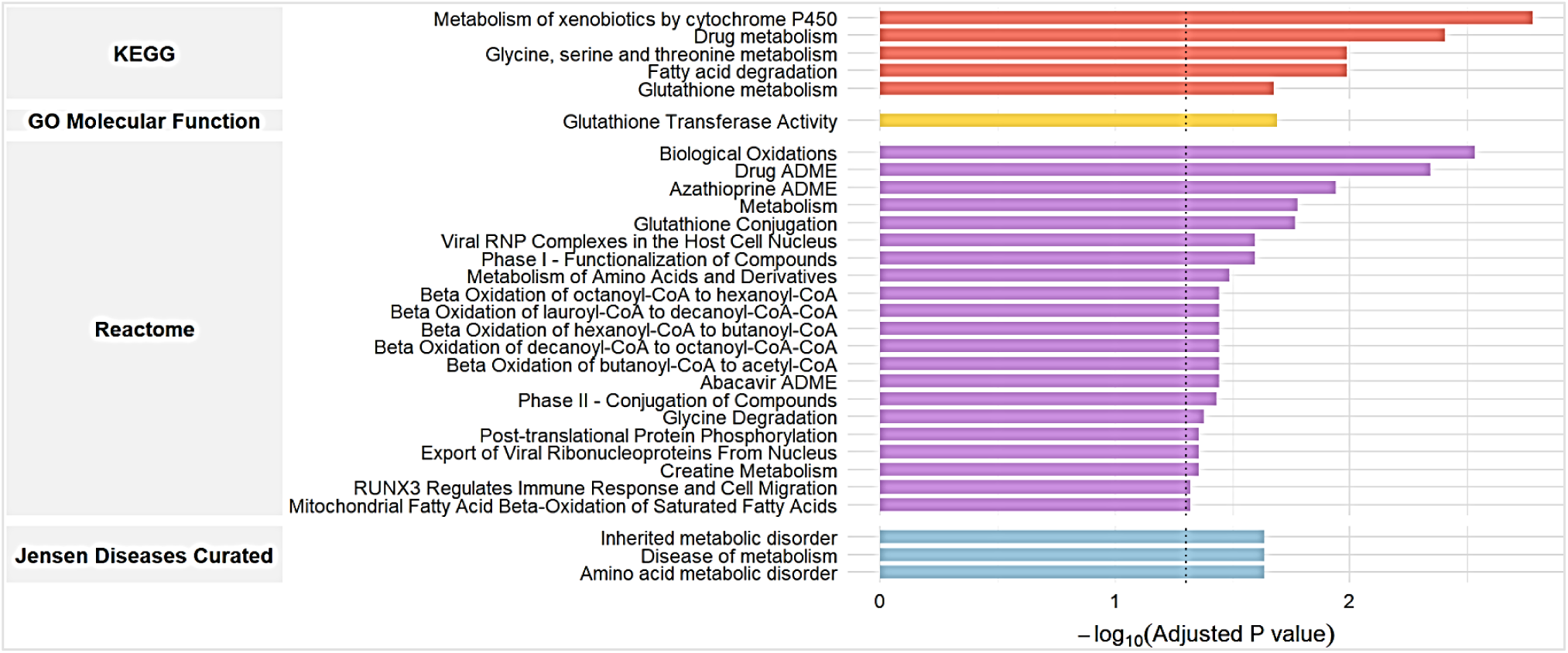
P**a**thway **analysis -** Pathway enrichment analysis of differentially expressed proteins between normal and abnormal endothelial function groups with unadjusted p value < 0.005. The bar plot shows significantly enriched pathways according to KEGG, Gene Ontology (GO) Molecular Function, Reactome, and Jensen Diseases Curated databases. The x-axis represents the - log₁₀(adjusted p value) for each pathway, with a higher value indicating stronger enrichment. Top pathways include drug metabolism and xenobiotic metabolism (KEGG), glutathione transferase activity (GO), biological oxidations (Reactome), and inherited or metabolic diseases (Jensen Diseases). The black dashed line denotes the adjusted p-value threshold of 0.05 for statistical significance.

Figure 3 shows that the KEGG database identified pathways related to drug metabolism, glutathione metabolism and fatty acid degradation as specifically dysregulated in participants with impaired CEF. The GO Molecular Function database identified pathways from the glutathione transferase activity pathway. The Reactome database identified pathways like fatty acid β-oxidation, glutathione conjugation and metabolism of amino acids or glycine degradation, as well as viral and immune mediated pathways like - viral RNP complexes and RUNX3 regulation of immune response. The Jensen Disease database identified pathways related to diseases of metabolism.

Collectively these pathway analyses indicated that endothelial dysfunction is associated with dysregulation of biological pathways related to metabolism, in particular glutathione metabolism/detoxification and fatty acid oxidation.

## DISCUSSION

In this study combining *in vivo* coronary MRI and high-throughput serum proteomics, we found that CEF impairment is highly prevalent in PWH when compared to PWoH despite viral suppression and comparable traditional cardiovascular risk profiles.

Nearly two-thirds of the PWH studied, demonstrated impaired coronary vasoreactivity, whereas the majority of PWoH exhibited preserved endothelial function. Despite this marked physiological difference, global proteomic profiles between PWH and controls were similar. However, when participants were stratified by endothelial function, distinct proteomic signatures emerged, highlighting enrichment in metabolic pathways related to glutathione metabolism and fatty acid oxidation. These findings implicate metabolic and oxidative dysregulation in endothelial dysfunction and suggest that HIV-status driven alteration in metabolic and oxidative processes might be central drivers of impaired CEF and cardiovascular disease in virally suppressed, treated HIV patients.

This study fills a critical knowledge gap by directly linking a quantitative vascular phenotype (CEF measured by MRI) with comprehensive proteomic profiling.

Whereas prior serum proteomic studies have compared PWH and controls to investigate remodeling of cardiac chambers (22), or extracardiac organ damage (23), our approach integrates functional vascular data, allowing us to detect molecular signatures truly associated with early-stage vascular damage. These results support a paradigm in which residual cardiovascular risk in treated HIV stems from persistent metabolic and oxidative imbalances that compromise local vascular homeostasis.

### Relation to Prior Studies and Novel Insights

Our findings extend prior work demonstrating that HIV-related endothelial dysfunction persists despite viral suppression and conventional risk factor control (14,24). Previous studies have largely focused on systemic inflammation and immune activation as mediators of disease in PWH (22); however, accumulating data points to a pivotal role for oxidative stress and metabolic injury (25). Experimental studies in HIV-1 transgenic rats have shown that chronic viral protein expression depletes vascular nitric oxide (NO) and reduces antioxidant capacity, which are effects that were reversed by glutathione restoration (26). Our findings suggest that these observations might also be relevant to humans.

HIV infection causes increased ROS production in patients along with a decrease in glutathione levels due to inhibition of glutathione synthesis. Glutathione is a major endogenous antioxidant, and its diminishing supplies can disturb antioxidant defense systems (27). This cascade can lead to eNOS uncoupling, which diminishes nitric oxide production and contributes to endothelial dysfunction. Sufficient glutathione stores are essential to maintain redox equilibrium, preserve vascular homeostasis, and maintain normal immune cell function (28). In our study, we observe enrichment of pathways associated with glutathione transferase. The enrichment of glutathione metabolism and conjugation pathways in HIV participants with impaired CEF suggests a sustained adaptive antioxidant response to oxidative stress. This likely represents a compensatory mechanism that is insufficient to restore endothelial redox homeostasis, contributing to nitric oxide depletion and impaired vasoreactivity.

Mitochondrial dysfunction and alteration of oxidative processes are a well-documented feature of chronic HIV infection, driven by viral proteins (e.g. Tat, Vpr) and lasting ART exposure, which disrupt mitochondrial DNA integrity and impair oxidative phosphorylation (25,29). Impaired mitochondrial respiration promotes electron leakage, ROS generation, nitric oxide depletion through eNOS uncoupling, and reduced glutathione that further weakens antioxidant defenses. This creates a cycle of oxidative stress and endothelial injury (25). In addition, HIV-related metabolic reprogramming, characterized by elevated glycolysis and impaired β-oxidation, results in accumulation of acylcarnitines and sustained immune and metabolic stress that persists despite viral suppression, thus contributing to vascular dysfunction and atherosclerotic progression (30).

We found that proteins involved in fatty acid degradation and glutathione metabolism were dysregulated in participants with abnormal CEF. These alterations point to metabolic disturbance involving glutathione pathways and oxidative stress as fundamental drivers of vascular risk in the treated HIV population. Emerging mechanistic studies suggest that SGLT2 inhibitors and GLP-1 receptor agonists favorably modulate mitochondrial metabolism, fatty acid use and oxidative balance in cardiac and metabolic tissues. This may represent therapeutic strategies to target similar pathways of metabolic or oxidative stress in vascular disease (31,32). Taken together, the integration of vascular imaging and proteomics therefore offers a mechanistic framework to guide therapeutic targeting of residual cardiovascular risk in PWH.

### Limitations

This study has several limitations. First, the sample size was modest and derived from a single-center cohort, which may limit statistical power for detecting subtle proteomic differences by HIV status. Second, the cross-sectional design precludes causal inference between the identified proteomic pathways and endothelial dysfunction. Third, although we used a high-throughput proteomic platform (SomaScan 7K), the coverage of low-abundance signaling proteins remains incomplete. Finally, although stress coronary MRI offers a robust physiological assessment of endothelial function, incorporating complementary measures such as peripheral vascular testing or other tissue redox biomarkers would further strengthen the mechanistic interpretation.

### Conclusions

In summary, PWH demonstrate marked coronary endothelial dysfunction despite viral suppression and minimal systemic proteomic differences. Proteomic signatures associated with impaired CEF suggest that metabolism and fatty acid oxidation might be important contributors to endothelial dysfunction rather than HIV-specific inflammatory processes. These findings support a mechanistic model in which oxidative and metabolic stress drive residual vascular risk in HIV. Targeting these pathways may offer a promising strategy to improve vascular health in the growing population of people aging with HIV.

## Funding

This work is supported by NIH grants NIH/NHLBI R35HL172680 (AGH) and NIH/NHLBI T32HL007227 (SB).

## Disclosures

Luigi Adamo is a consultant for Kiniksa pharmaceuticals and Novo Nordisk and a founder of i-Cordis, LLC.

## Notes

### Competing Interest Statement

The authors have declared no competing interest.

