## Supplementary Figure 1 for "Metabolic and Redox Pathway Dysregulation in HIV-Associated Coronary Endothelial Dysfunction"

**A:** **B:**

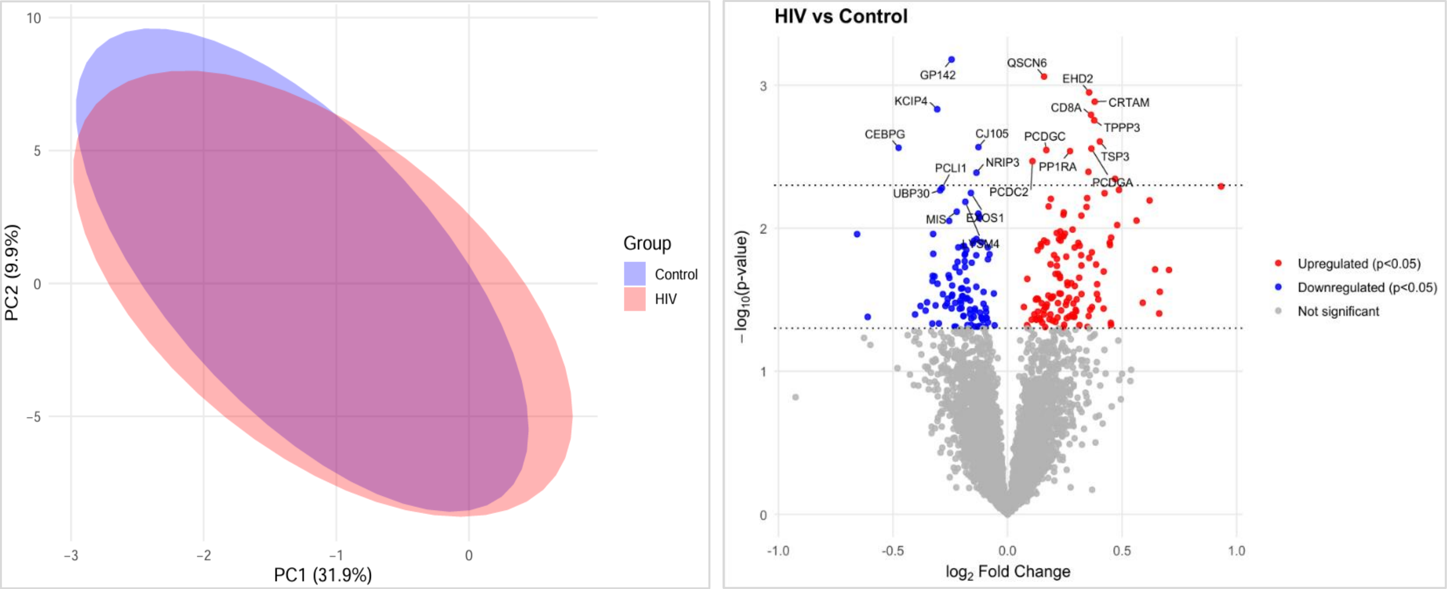


**Supplementary Figure 1:** Serum proteomic analysis of patients with and without HIV. **A)** Principal component analysis (PCA) of study participants by HIV status. The plot shows the distribution of control (purple) and HIV (peach) groups along the first two principal components. There is a substantial overlap between groups, indicating only partial separation in their global expression profiles. **B)** The volcano plot represents the differential protein expression between participants with HIV and without HIV (Control). The volcano plot displays $\log_2$ Fold Change (x-axis) versus $-\log_{10}$(p-value) (y-axis) for each protein. Proteins with an unadjusted p-value of less than 0.05 are labelled red (upregulated) or blue (downregulated). The top 10 upregulated and downregulated proteins are labeled.
