## Supplementary Figure 2 for "Metabolic and Redox Pathway Dysregulation in HIV-Associated Coronary Endothelial Dysfunction"

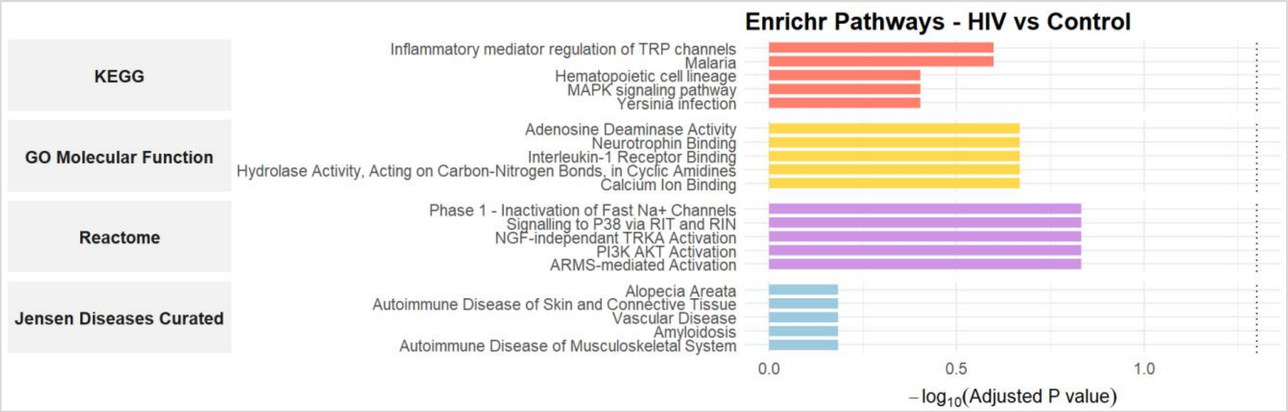


**Supplementary Figure 2:** Pathway analysis - Pathway enrichment analysis of differentially expressed proteins between normal and abnormal endothelial function groups with unadjusted p value < 0.005. The bar plot shows the top 5 enriched pathways according to KEGG, Gene Ontology (GO) Molecular Function, Reactome, and Jensen Diseases Curated databases. The x-axis represents the $-\log_{10}$(adjusted p value) for each pathway, with a higher value indicating stronger enrichment. Not a single pathway in any of these databases had an adjusted p-value of less than 0.05 which indicates that no pathways were enriched significantly. The black dashed line denotes the adjusted p-value threshold of 0.05.
