## Supplementary Table 1 for "Metabolic and Redox Pathway Dysregulation in HIV-Associated Coronary Endothelial Dysfunction"

**Supplementary Table 1: Additional characteristics of patients living with HIV**

| **Measure** | **Mean ± SD or %** |
|---|---|
| **CD4 count, cells/μL** | 693.1 ± 375.7 |
| **CD4 nadir, cells/μL** | 336.0 ± 311.5 |
| **CD4/CD8 ratio** | 1.78 ± 2.25 |
| **On ART** | 98% |
| **Protease inhibitor use** | 16% |
| **NNRTI use** | 18% |
| **NRTI use** | 73% |
| **INSTI use** | 82% |

**Abbreviations: NNRTI = non-nucleoside reverse transcriptase inhibitor; NRTI = nucleoside reverse transcriptase inhibitor; INSTI = integrase strand transfer inhibitor.**
