## Supplementary Table 2 for "Metabolic and Redox Pathway Dysregulation in HIV-Associated Coronary Endothelial Dysfunction"

**Supplementary Table 2. Baseline Characteristics by Coronary Endothelial Function (CEF) Status**

| Variable | | HIV- (n = 29) | HIV+ (n = 45) | p-value |
|---|---|---|---|---|
| **Age, years (mean ± SD)** | | 52.4 ± 19.4 | 56.9 ± 12.4 | **0.23** |
| **Female sex, %** | | 34 | 49 | **0.22** |
| **Race, %** | **Black** | 45 | 62 | **0.14** |
| | **White** | 34 | 31 | **0.76** |
| | **Asian** | 10 | 2 | **0.13** |
| **Ethnicity, %** | **Hispanic** | 10 | 7 | **0.62** |
| | **Non-Hispanic** | 90 | 91 | |
| **BMI, kg/m² (mean ± SD)** | | 27.0 ± 7.6 | 29.3 ± 4.8 | **0.12** |
| **Hypertension, %** | | 71 | 62 | **0.56** |
| **Diabetes, %** | | 14 | 7 | **0.35** |
| **Smoking history, %** | | 25 | 40 | **0.20** |
| **Smoking, packs/day (mean ± SD)** | | 0.79 ± 0.62 | 1.02 ± 0.79 | **0.49** |

| | | | | |
|---|---|---|---|---|
| **History substance abuse, %** | | 26 | 43 | **0.15** |
| **Total cholesterol, mg/dL (mean ± SD)** | | 176.0 ± 37.3 | 183.6 ± 42.3 | **0.45** |
| **LDL cholesterol, mg/ (mean ± SD)** | | 96.4 ± 32.4 | 108.1 ± 39.2 | **0.20** |
| **HDL cholesterol, mg/dL (mean ± SD)** | | 60.7 ± 18.9 | 53.8 ± 15.4 | **0.096** |
| **Triglycerides, mg/dL (mean ± SD)** | | 98.0 ± 71.3 | 115.7 ± 52.6 | **0.23** |
| **Abnormal coronary endothelial function, %** | | 10 | 67 | **<0.001** |
| **% Change in Coronary Cross-Sectional Area (mean ± SD)** | | 4.28 ± 16.99 | −2.99 ± 10.99 | **0.037** |
